# LVING Reveals the Intracellular Structure of Cell Growth

**DOI:** 10.1101/2023.09.08.553132

**Authors:** Soorya Pradeep, Thomas A. Zangle

## Abstract

The continuous balance of growth and degradation inside cells maintains homeostasis. Disturbance of the balance by internal or external factors cause state of disease. Effective disease treatments seek to restore this balance. Here, we present a method based on quantitative phase imaging (QPI) based measurements of cell mass and the velocity of mass transport to quantify the balance of growth and degradation within intracellular control volumes. The result, which we call Lagrangian velocimetry for intracellular net growth (LVING), provides high resolution maps of intracellular biomass production and degradation. We use LVING to quantify the growth in different regions of the cell during phases of the cell cycle. LVING can also be used to quantitatively compare the effect of range of chemotherapy drug doses on subcellular growth processes. Finally, we applied LVING to characterize the effect of autophagy on the growth machinery inside cells. Overall, LVING reveals both the structure and distribution of basal growth within cells, as well as the disruptions to this structure that occur during alterations in cell state.

## INTRODUCTION

Maintaining the balance of biomass production and degradation defines the health of cells and the overall health of the organism^1^. For example, proteins are generated continuously enabling cell growth while improperly folded proteins are degraded which aids in degenerative diseases^2^. Similarly, dysregulation of any cellular process can cause or contribute to disease, including diabetes, cancer, and organ myopathy, like cardiac myopathy^3,4^. Studying these diseases and their treatments requires measuring growth and how therapies act to restore the balance of growth.

One widely used approach to measure cell growth is to incorporate labels that can be tracked over time. For instance, DNA fiber autoradiography^5^, fluorescent Feulgen assay^6^, or flow microfluorography quantify DNA replication^7^. These techniques are highly accurate but require the introduction of radioactive or toxic foreign materials that may interfere with normal cell function. Simple cell proliferation assays can be a non-invasive alternative, as this process simply relies on the periodic counting of cells^8^. However, proliferation assays typically measure only population average growth, and do not give details on cellular processes. We can overcome these disadvantages using techniques such as microchannel resonators^9^, deep UV imaging^10,11^, Raman imaging^12^, and quantitative phase imaging (QPI)^13^. Microchannel resonators are precise to the femtogram level^14^, very high sensitivity relative to the ∼100 pg mass^9^ of a typical cell, but do not quantify growth within single cells. UV and Raman microscopy can be used to measure the protein, nucleic acid, and lipid content of cells. Of these, UV microscopy has high resolution due to the low wavelength ultraviolet light^15^, but can be used only for short term measurements as extended UV light exposure triggers cell death. Normalized Raman microscopy offers high-resolution measurement of protein, lipid, and nucleic acid content with modifications that eliminate error from the scattering of light^12^ but is still less widely used due to complex optics and cost. QPI has been used to precisely measure cell dry mass with multiple commercial options on the market^13^.

QPI is a non-invasive, label-free technique that measures the phase shift of visible light as it passes through a cell. This phase shift is proportional to the dry biomass distribution inside of cells^16^. The rate of increase of cell biomass over time indicates the cell growth rate^17^, and the impact of cancer drugs on decreasing growth rate can also be measured via QPI^18^. Improvements in post-processing of QPI data to increase the precision of dry mass measurements reveals significant oscillations in cell mass during the cell cycle, reflecting a balance of growth and degradation^19^. However, while these and other QPI studies reveal the details of growth to the single cell level^20^, this approach is not capable of quantifying the distribution of growth within these individual cells. Deep learning methods have been applied to segment nucleus from cytoplasm to quantify overall growth of these compartments^21^. In this work we take the next step towards spatially resolved measurements of growth within cells.

Here, we develop Lagrangian velocimetry for intracellular net growth (LVING) to quantify sub-cellular cell growth from QPI data. Both biomass production/destruction and motion of mass can cause an apparent change in biomass at each location within the cell. We use the principles of particle image velocimetry (PIV) to measure the intracellular biomass velocity field from QPI data^22^. LVING then discretizes the cells into small volumes and estimates the rate of change of biomass in each volume after correcting for cell motion. We validate LVING by measuring the localization of biomass generation in cells during the cell cycle. As expected, we see growth in the perinuclear region of cells in the G1 phase, intensified growth inside the nucleus in S phase and growth in the peri-nuclear region as the cell goes through G2 phase. We also demonstrate LVING for measuring the effect of different protein or DNA-synthesis inhibitors on the patterns of mass accumulation within cells. Finally, we apply LVING to study the intracellular impacts of autophagy and observe a decrease in cytoplasmic growth.

## RESULTS

### LIVING maps growth distribution in single live cells

We developed Lagrangian velocimetry for intracellular net growth (LVING) to measure the growth in discretized volumes inside of single cells. First, we use QPI based on quadri-wave lateral shearing interferometry (QWLSI)^23^ to image intracellular distribution of phase shifts (**figure S1**)-with high accuracy and precision (**figure S2**) This phase shift data is related to mass using the specific refractive increment^13,24^ resulting in images of the distribution of dry biomass (mass excluding water) within single cells (**figure 1a**). Sequences of biomass distributions can then be used to measure the overall change in biomass over time within cells (**figure 1b**). This change in biomass at any given (Eulerian) location (control volume) within the cell is due to both motion and biosynthesis or degradation of mass. We can express this in terms of a mass balance:

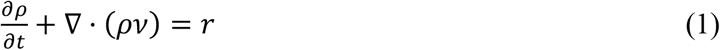

which states that the change in mass change over time, 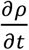, at any location within the cell is due to both the net mass flux in and out of that location, ∇ · (*ρ ϑ*), and the net mass generation, *r*. We measure the biomass flux using intracellular velocity measurements made with quantitative phase velocimetry (QPV)^22^ (**figure 1c**). We then translate the Eulerian measurements of the rate of change of mass (**figure 1d**) to a Lagrangian frame of reference that tracks control volumes within the cells in space and over time (**supplementary derivation)**. The resulting change in mass in each Lagrangian control volume is due to the generation or degradation of mass. We then measure the mass accumulation rate of each individual control volume (**figure 1e**), resulting in a map of growth rates, rate of change of mass with time, within 0.7 μm^2^ regions in the cell (**figure 1f**). Results for artificially moved fixed cells show the expected result of no growth (**figure S3**). Normalizing the sum of growth rates of volumes over an area inside cell by the total mass in the region gives the specific growth rate of that region of the cell. As expected, the total specific growth rate in all volumes inside RPE and MCF7 obtained from LVING matches the overall growth rate computed with QPI and whole cell segmentation (**figures S3c**,**f** and **S4**).

**Figure 1.**
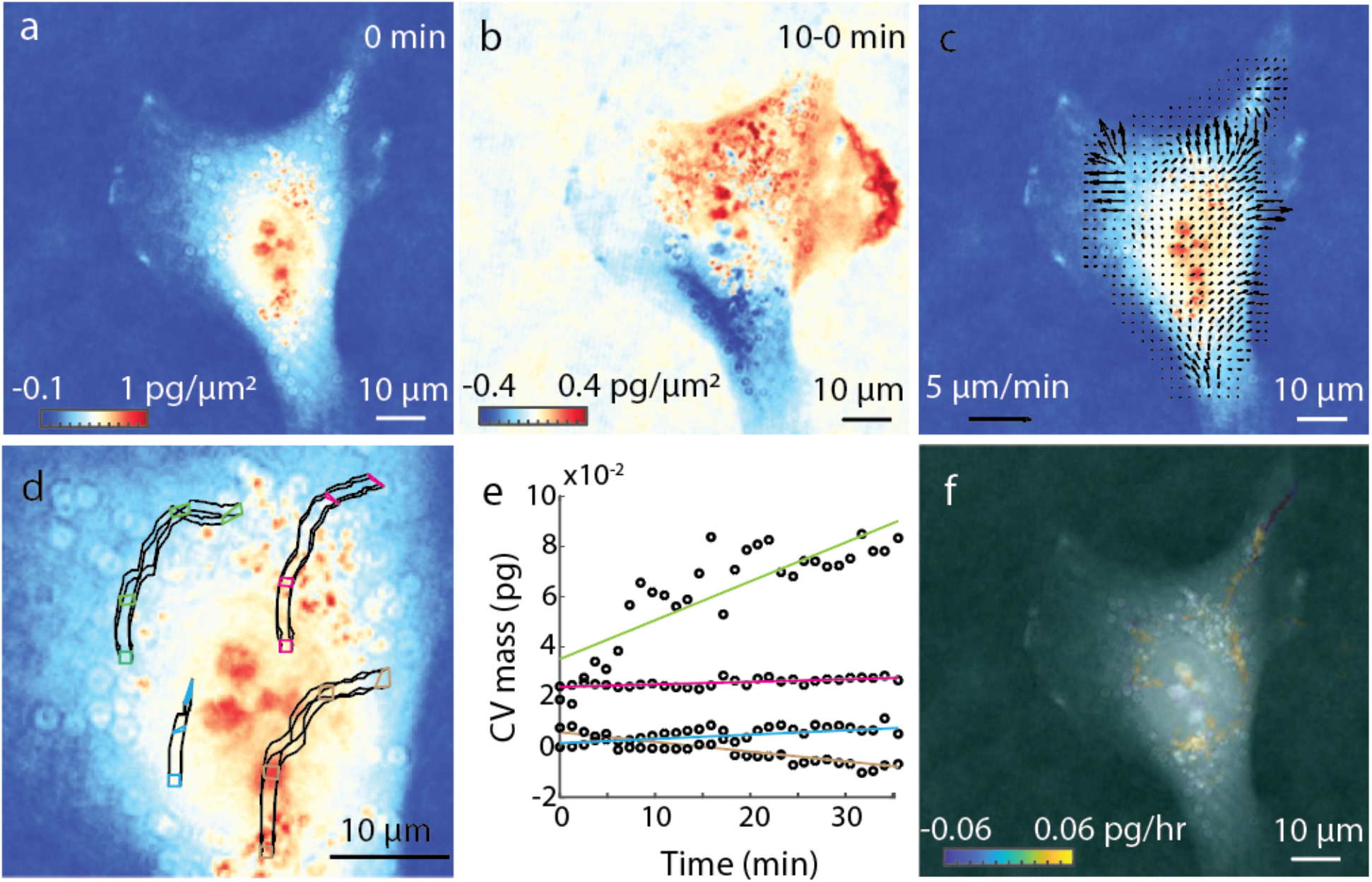
Lagrangian Velocimetry Intracellular Net Growth (LVING) uses quantitative phase images plus intracellular velocimetry to measure growth inside cells. (a) We use quantitative phase imaging (QPI) based on wavefront sensing to measure the phase shift of light passing through a sample (here: RPE cells), which is proportional to the dry biomass distribution within cells. (b) The difference between QPI mass distributions over 10 min shows motion of mass. (c) Intracellular velocimetry computes the rate and direction of mass transport. (d) Intracellular velocity vectors are used to track the movement and deformation of control volumes over time (4 shown here over 30 min). (e) the slope of the rate of change of mass of each control volume over time is them computed to find the growth rate within each control volume. (f) Computed growth map in the cell over 30 min showing localized growth within the cell as distinct puncta.

### LVING maps intracellular biomass fixation dynamics during cell cycle

We performed LVING on RPE and MCF7 cells to visualize the localization of growth during different phases of the cell cycle in overlapping 2-hour windows (**figure 2** and **S5, movie M1** and **M2**). LVING growth maps in RPE cells show growth in both cytoplasmic and nuclear regions of the cells concentrated within regions of high mass density (**figure 2a,d,e,h**). We identified the cell cycle phase using FUCCI markers (**figure 2b,c,f,g**). We then segmented nuclei from the cell cytoplasm using the FUCCI marker and tracked this location over time (**figure S6**). We calculated the specific growth rate in the nucleus and cytoplasm from the average growth rate at all control volumes inside the nuclear and cytoplasmic regions from the LVING growth map (**figure 2i**). The specific growth rate in the cell cytoplasm is uniform over the cell cycle, whereas the cell nuclear specific growth rate is negligible in G1 and S phase and higher in the G2.

**Figure 2.**
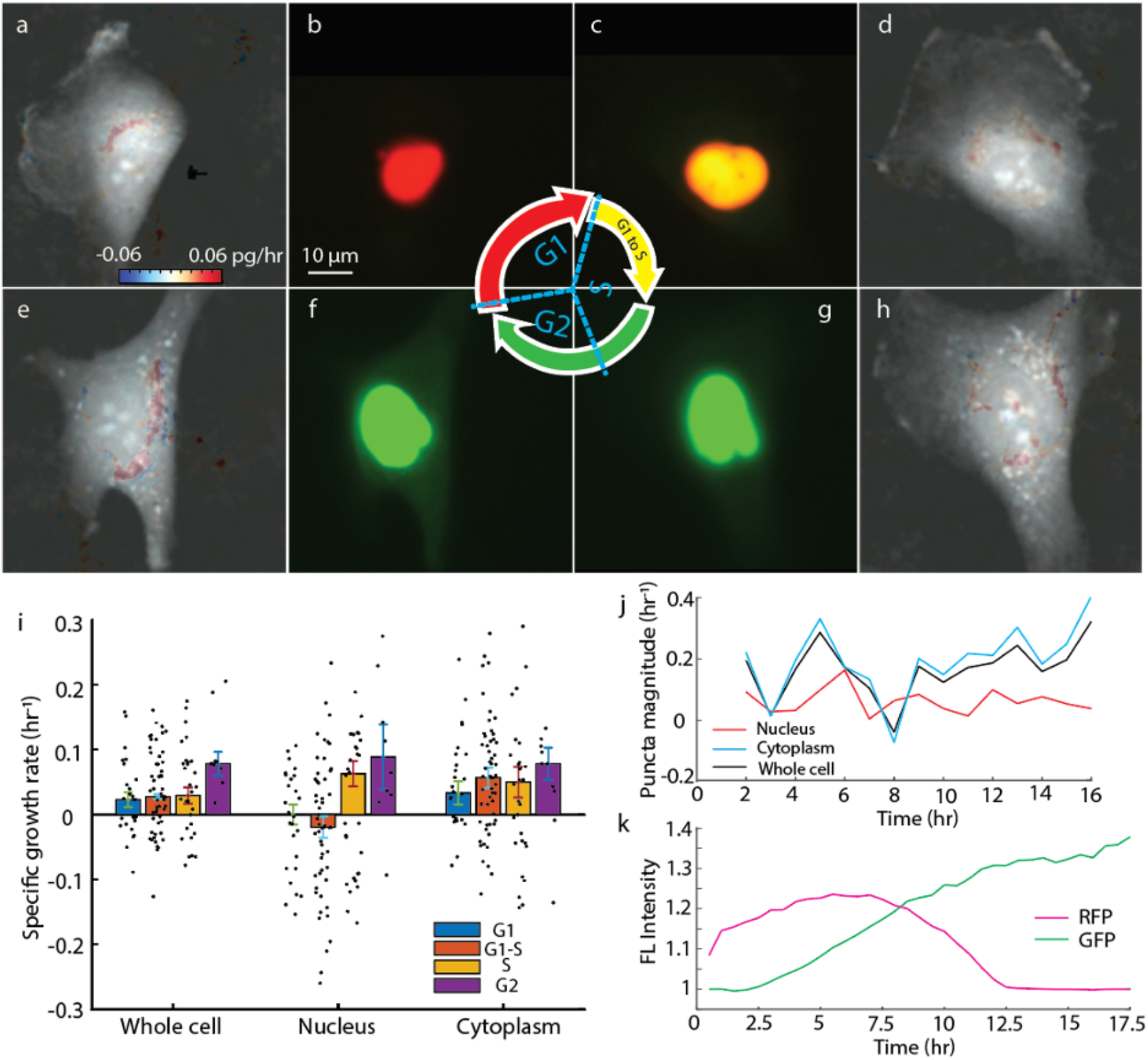
Quantitative intracellular growth map measured with LVING reveals growth puncta in nucleus and cytoplasm of RPE cells. (a) to (c) shows intracellular growth maps for a single RPE cell tracked through the G1, S and G2 phases of the cell cycle. The red marker indicates the segmented nucleus, the scalebar indicates 10 μm, and the colorbar indicates the growth rate in control volumes in pg/h. (d) to (f) shows the FUCCI marker corresponding to the cells images in (a) to (c) red nucleus tagging indicates cell is in G1 phase, yellow is S phase and green is G2 phase of cell cycle. (g) Specific growth rate of RPE cells averaged over the whole cell, inside the segmented nucleus and within the segmented cytoplasm during G1, S and G2 phases of cell cycle.

The growth pattern in MCF7 cells shows increased growth immediately around the nucleus, in the perinuclear region (**figure S5**) as expected for localization of protein production. Nuclear growth puncta are visible persisting through the G2 phase of the cell cycle. In addition to observations of perinuclear rings of biomass synthesis (e.g. **figure 2e, S5e**) LVING data also enables observation of localized growth puncta and their evolution through the cell cycle (**figure 3**).

**Figure 3.**
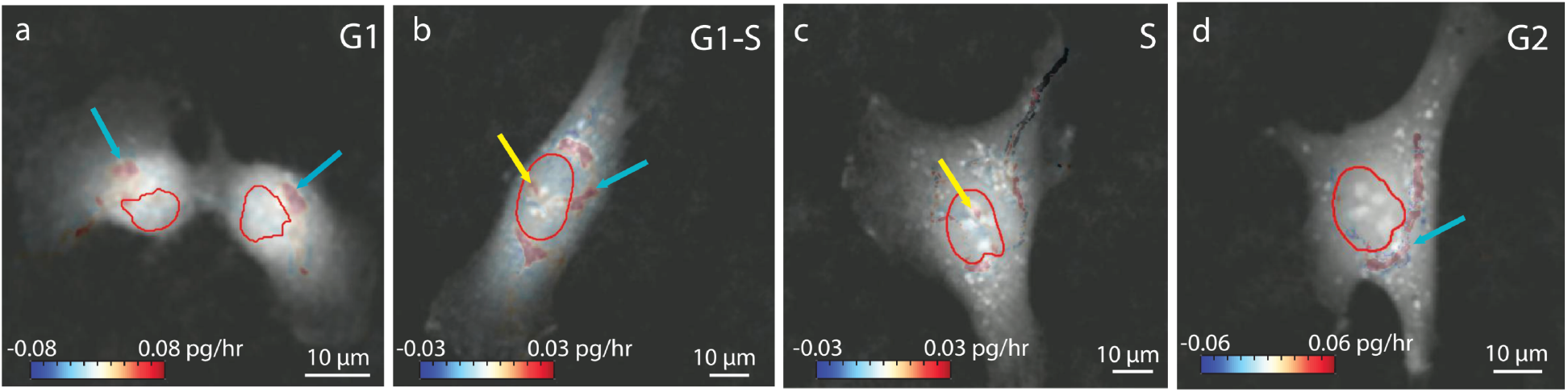
Localization of growth puncta in nucleus and cytoplasm of RPE cells during phases of cell cycle. (a) Growth puncta localized in cytoplasm during G1 phase (blue arrow). (b) Growth puncta starts appearing in nucleus during the transition to S phase from G1 phase. (c) Growth puncta increases in nucleus in S phase. (d) Additional regions of growth visible in cytoplasm during G2 phase as the cell prepares to divide.

Doxorubicin is a topoisomerase II drug which inhibits the DNA synthesis inside the nucleus^25^. The inhibitory concentration of doxorubicin in RPE cells is around 0.1 μM for longer period of treatment^26^, so we chose to test doses at lower and higher concentrations over 6 h. We observed the effect of doxorubicin at 0.01, and 0.2 μM doses on RPE cells. The intracellular growth map of RPE cells treated with 0.2 μM doxorubicin show elimination of growth puncta within the nucleus (**figure 4**), though puncta are retained at 0.01 μM (**figure S7**). Similarly, cells treated with homoharringtonine, a protein synthesis inhibitor^27^, show a reduction in cytoplasmic growth rates with increasing dose (**figure S8**).

**Figure 4.**
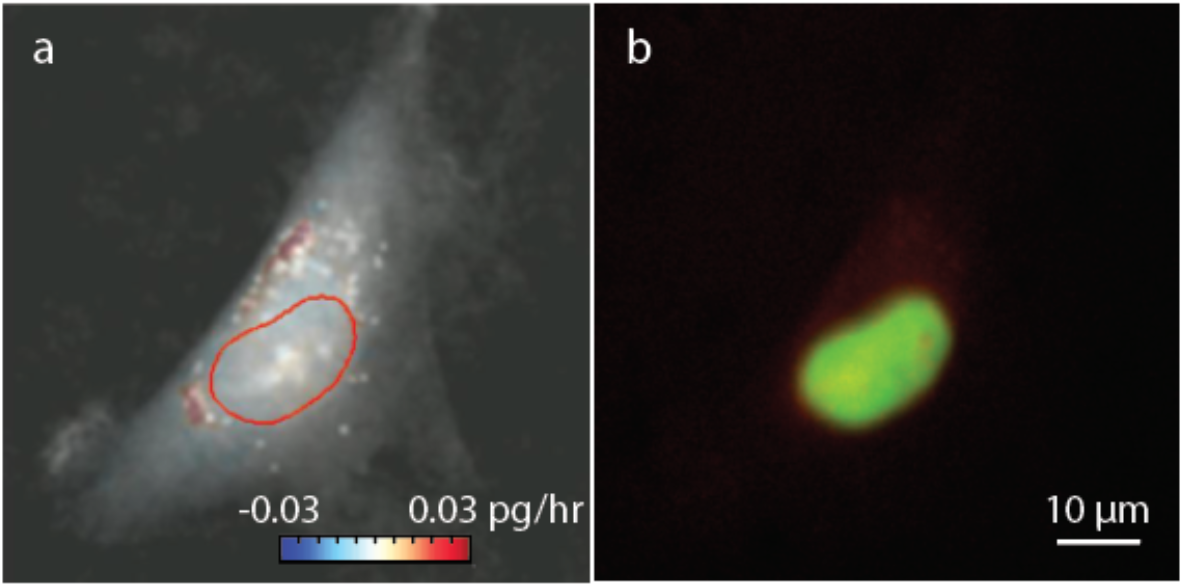
Effect of doxorubicin, a specific inhibitor of topoisomerase II, which is required for DNA synthesis, on RPE cells. (a) Intracellular growth map of RPE cell treated with 0.2 μM doxorubicin. Colorbar indicates growth rate in pg/hr. (b) Corresponding FUCCI marker indicates the cell shown in (a) is in the S/G2 phase of cell cycle.

### Gradual shut-down of growth in autophagic cells

Cell death happens in numerous modes, including autophagy, necrosis, entosis, and apoptosis^28^. Autophagy is a mode of slow death of cells in which cells degrade their own constituents by transporting them in lysosomes fusing into autophagosomes in the cytoplasm, where they are digested^29^ and is induced in cells through nutrient starvation^30^. We induced autophagy in RPE and MCF7 cells in this work by complete nutrition deprivation using Earl’s balanced salt solution (EBSS) starvation media^31^. Autophagy in the cells was confirmed by Western blot of MCF7 and RPE cells undergoing autophagy (**figure S9**). During autophagy, the materials to be degraded are tagged with LC3-1 protein and are encapsulated in the lysosomes. The LC3-1 tagged constituents are called LC3-2. The LC3-2 is transported in the lysosomes which fuse with the autophagosomes in the cytoplasm, transferring them into the autophagosomes. The pH in the autophagosome is low facilitating the degradation. We cannot observe the quantity of materials which are already degraded in the Western blots. In order to observe the material degraded during autophagy, we inhibit the last step of LC3-2 degradation by using chloroquine. Chloroquine increases the pH in the autophagosomes in the cells disrupting the degradation and thus showing an increased intensity of LC3-2 band in Western blot, compared to just EBSS treated cells undergoing autophagy (**figure S9a** and **c**).

We then observed the inhibition of RPE cell growth using QPI data, which captured a gradual decrease in the cytoplasm biomass production over time in MCF7 (**figure 5**) and RPE cells (**figure S10**). We tagged the MCF7 cells with a lysosome marker to observe the accumulation of the autophagosomes. The cells accumulate the materials to be degraded in the autophagosomes during autophagy. We observed an increase in lysosome concentration in MCF7 cells over time due to the progress of cell autophagy (**figure 5e**). Growth in EBSS treated cells reduces over time, eventually shutting down completely (**Figure 5d, S10d**).

**Figure 5.**
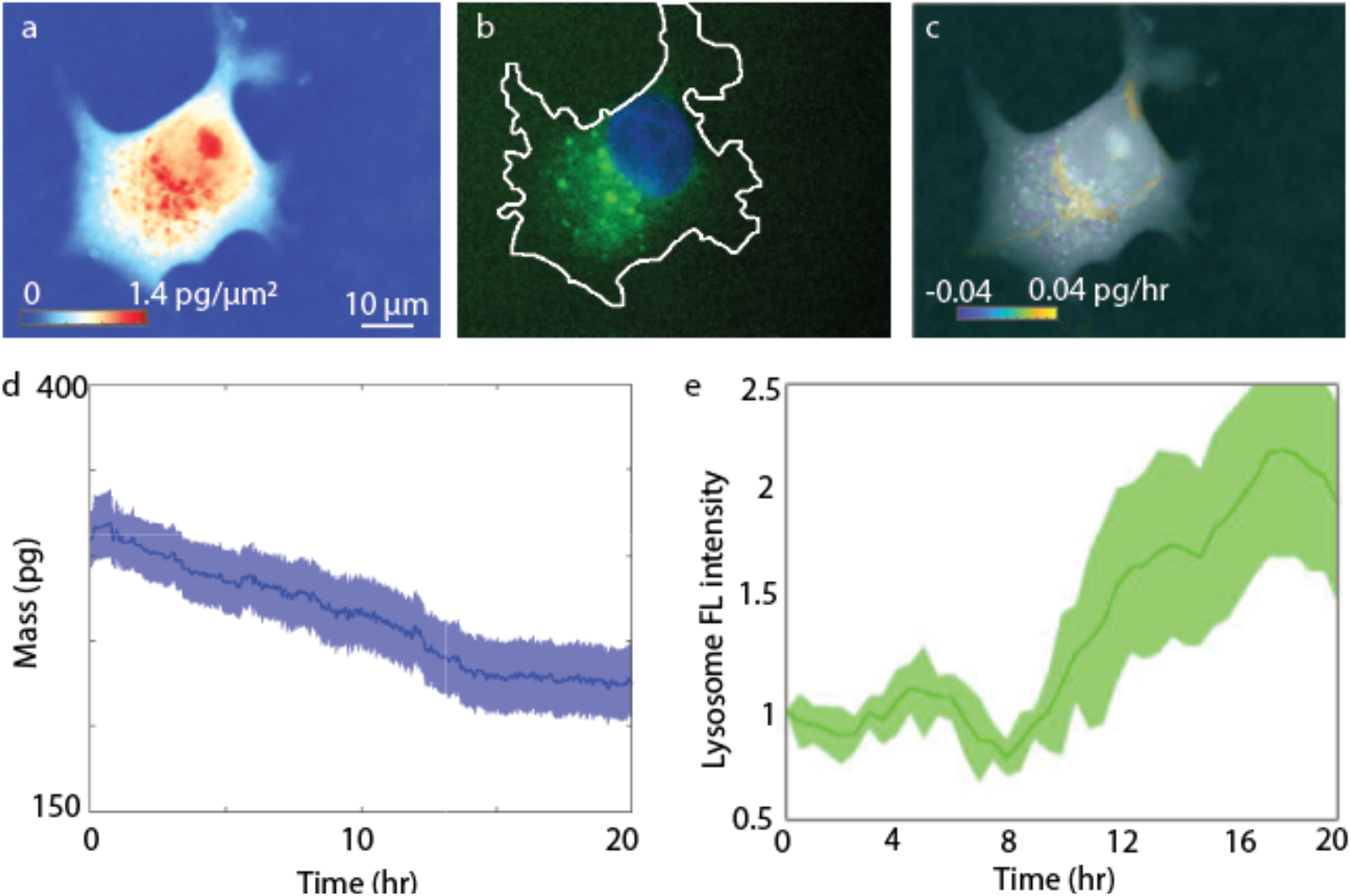
LVING provides a tool to visualize the process of autophagy in MCF7 cells. (a) QPI image of MCF7 cell undergoing autophagy. Scalebar indicates 10 μm and colorbar indicates dry mass in pg/μm2. (b) GFP tagged LC3 marker shows lysosome accumulation in cell cytoplasm due to cell autophagy. (c) Growth rate map from LVING at the start of the experiment shows growth still prevalent at perinuclear region, but negative degradation puncta in the surrounding regions of the cytoplasm. (d) QPI whole cell dry mass vs time data shows (n=21) biomass degradation in cells due to autophagy. (e) Accumulation of lysosome marker intensity from the same cells, normalized by initial intensity at start of imaging, shows change in the fluorescence intensity over experiment time as an indication of progress of autophagy.

## METHODS

### Cell culture and FUCCI tagging

mKO2-hCdt1- and mAG-hGem-tagged FUCCI RPE-1 cells were acquired from the lab of Bruce Edgar (University of Utah) and were cultured in DMEM with 10% fetal bovine serum (FBS) and 5% penicillin-streptomycin (penicillin-streptomycin was washed out prior to imaging). MCF7 cells were donated by the lab of Bryan Welm (University of Utah) and were cultured in DMEM 1ith 10% FBS. MCF7 cells were transiently transfected with Premo FUCCI cell cycle sensor tags (Thermofisher, USA). MCF7 cells required additional treatment with BacMam enhancer (Fisher Scientific, USA) to express FUCCI following manufacturer instructions.

### Drug treatment

Doxorubicin (Fisher, USA),was prepared in DMSO and diluted to the target final concentration in complete cell culture media with no antibiotic. Then cell culture media was aspirated from the dish and is replaced with the drug containing media and the dish was returned to the incubator for a few minutes to stabilize the temperature before moving to the microscope stage for imaging.

### Induction and confirmation of autophagy

2000 to 4000 cells were plated in 200 μL of media in a 96 well plate and incubated for 24 h before mixing each of Premo Autophagy Sensors (LC3B-FP, ThermoFisher, USA) BacMam 2.0, LC3B-FP (Component A) and LC3B(G120A)-FP (Component B) (ThermoFisher, USA) by inversion to ensure a homogenous solution. 1 μL of each of the LC3B reagent was added to the cells for a target multiplicity of infection (MOI) of 3 and cells were incubated overnight (≥16 h). 4 h after adding the LC3B reagents, chloroquine (Signa-Aldrich) was added to make up a concentration of 30 to 100 μM in the wells and incubated for 12 to 16 h. After incubation, cells were stained with 10 μg/mL DAPI (Fisher Scientific) for 5 to 15 minutes before washing with fresh media and being allowed to attach to the dish. Finally, the nutrient rich media was removed and cells were washed 3x with Earle’s balanced salt solution (EBSS) to induce starvation, leaving a total of 1 mL EBSS starvation media in the 35 mm ibidi dish (ibidi, Germany) used for 24 h live cell imaging.

For Western blot confirmation of the EBSS protocol, cells were plated in 3 ml of medium per well in a six-well plate to obtain 80%–90% confluency by 48 h. At 24 h after plating, culture medium was removed from the well and replaced with fresh media in control well 1 and well 3, with EBSS added to wells 2 and 4 before an additional 24 h incubation. 2 h before harvesting chloroquine was added at a final concentration of 60 μM to wells 3 and 4. At harvesting, the plate was placed on ice, washed 2x with ice-cold PBS, then 50–100 μL pre-cooled lysis buffer was added and cells were collected with a cell scraper and tranfserred into a pre-cooled 1.5 ml Eppendorf tube. Cells were sonicated for 10–15 sec to complete lysis, then centrifuged at 12,000 rpm for 10 min at 4°C. The supernatant was transferred to a new tube and all were adjusted to equal protein concentrations, then mixed with loading buffer (final 1x concentration) and boiled at 100°C for 5 min in a water bath, put on ice for 1–2 min, then spun briefly to bring all solution to the bottom of the tube. Pre-cast Novex gels (Thermofisher, USA) were rinsed with water assembled into the gel tank (Thermofisher, USA). Equal amounts of protein (10–20 μg) was loaded into the wells of the SDS-PAGE gel and run at 100 V until the dye reaches 1/4 inch from the bottom of the gel. PVDF membrane was soaked in methanol for 1–2 min, incubated in ice cold transfer buffer for 5 min before transfer. The gel was equilibrated in ice cold transfer buffer (Novex, Thermofisher, USA) for 10 min before transfer at 90 V for 70 min and blocking with 5% BSA blocking buffer while rocking for 1 h at room temperature. The membrane was cut at 25 kDa to separate actin and LC3 bands and primary antibody in 5% BSA blocking buffer (anti-LC3B, rabbit, Sigma-Aldrich L7543 at 1:1000 and anti-actin, mouse, Sigma-Aldrich A2228 at 1:2000) was incubated overnight at 4°C. PVDF was washed 3x in TBST and incubated for 1 h with secondary antibody in 5% BSA blocking buffer at room temperature with rocking, before washing 3x with TBST and imaging with LICOR Odyssey CLX (Licor, USA) at 685 and 785 nm.

### Imaging and image processing

Interferograms were acquired using a SID4BIO-4MP (Phasics, France) camera attached to an Olympus IX83 inverted microscope (Olympus Corporation, Japan) in brightfield with a 100x, 1.3 NA oil-immersion objective illuminated with a red (623 nm) Thorlabs DC2200 LED (Thorlabs, USA). Interferograms were acquired at 1 min intervals. A 1.2x magnifier was used to match the Nyquist criteria for reconstructed phase pixels. Matlab (Mathworks, USA) and Micromanager open-source microscopy software^32^ were used for image acquisition and processing. A flipping mirror (IX3-RSPCA, Olympus Corporation, Japan) was used for alternate fluorescence and QPI imaging. Fluorescence images were acquired every 30 min using a Retiga R1 camera (Cairn Research Ltd, UK) X-Cite 120LED (Excelitas Technologies, USA) illumination source. An Olympus U-FBNA filter cube was used for the green mAG fluorophore and a Semrock mCherry-B-000 filter cube (IDEX health & science, USA) was used for imaging the red mKO2 fluorophore. 37°C and 5% CO2 conditions were maintained using an Okolab stage-top incubator (Okolab, Italy) and custom-built objective heating collar, temperature controlled by Thorlabs temperature controller (Thorlabs, USA). Cells were plated in ibidi μ-high treated dishes (ibidi, Germany) at 30% confluence. Four sets of 30 live cells were imaged every for 8 h for every imaging session. Raw interferograms were processed to QPI data using the Phasics SDK (Phasics, France). A refractive increment of 0.18 μm^3^/pg was assumed for cell material. QPV^22^ was used to compute intracellular velocity vectors prior to LVING calculations. Image processing for LVING was computed using code available at https://github.com/Zangle-Lab/LVING

## DISCUSSION

Here, we developed LVING, a tool to map and quantify biomass production inside cells. LVING uses Lagrangian tracking of dry mass inside discretized control volumes inside cells, tracking them as they move and deform, and as they change mass due to macromolecule production or degradation. LVING was used to quantify the subcellular growth inside RPE and MCF7 cells during the cell cycle. as well as the disruption of cell homeostasis by treatment by specific inhibitors and on induction of the cellular death pathway of autophagy. LVING, therefore, provides the first spatially resolved measurements of growth within cells, based on a label-free platform that can be broadly applied to other cell types and disease models.

In application to measurements of actively cycling cells, LVING reveals large perinuclear puncta and/or rings of biosynthetic activity. In contrast, the observed growth within the nucleus is generally confined to smaller puncta whose total integrated magnitude is typically close the measurement noise floor (e.g. **figure 2j**). DNA typically constitutes a small fraction of the cell mass (<1%), so this is expected given the relative mass of protein that must be replicated each cell cycle compared to total DNA replication.

Finally, we note that, though in cells there is no actual ‘creation’ of mass, LVING assumes that we can equate the growth we observe with QPI to an effective generation term that captures the condensation of mass within the cell. This creates an effectively higher local concentration of biological macromolecules and other constituents than in the surrounding media. Therefore, LVING effectively captures the fixation of biomass that accompanies cell growth (e.g. **figure 2**) or intracellular processes that otherwise cause condensation of mass (e.g. **figure 5**). LVING, therefore, has potential applications to study any intracellular process that disrupts or displaces mass within the cell.

## REFERENCES

1 Cooper, J. P. & Youle, R. J. Balancing cell growth and death. Curr Opin Cell Biol 24, 802–803, (2012).

2 Hartl, F. U. Protein Misfolding Diseases. Annual Review of Biochemistry 86, 21–26, (2017).

3 Lloyd, A. C. The regulation of cell size. Cell 154, 1194–1205, (2013).

4 Yang, X. & Xu, T. Molecular mechanism of size control in development and human diseases. Cell Res 21, 715–729, (2011).

5 Friedkin, M., Tilson, D. & Roberts, D. Studies of deoxyribonucleic acid biosynthesis in embryonic tissues with thymidine-C14. J Biol Chem 220, 627–637, (1956).

6 Böhm, N. & Sprenger, E. Fluorescence cytophotometry: a valuable method for the quantitative determination of nuclear feulgen-DNA. Histochemie 16, 100–118, (1968).

7 <Cell Culture for Biochemists by R.L.P. Adams (Eds.) (z-lib.org).pdf<.

8 O’Brien, N. A. Activated Phosphoinositide 3-Kinase/AKT Signaling Confers Resistance to Trastuzumab but not Lapatinib (vol 9, pg 1489, 2010). Molecular Cancer Therapeutics 10, 2211–2211, (2011).

9 Godin, M., Delgado, F. F., Son, S., Grover, W. H., Bryan, A. K., Tzur, A. et al. Using buoyant mass to measure the growth of single cells. Nat Methods 7, 387–390, (2010).

10 Cheung, M. C., Evans, J. G., McKenna, B. & Ehrlich, D. J. Deep ultraviolet mapping of intracellular protein and nucleic acid in femtograms per pixel. Cytometry A 79, 920–932, (2011).

11 Cheung, M. C., LaCroix, R., McKenna, B. K., Liu, L., Winkelman, J. & Ehrlich, D. J. Intracellular protein and nucleic acid measured in eight cell types using deep-ultraviolet mass mapping. Cytometry A 83, 540–551, (2013).

12 Oh, S., Lee, C., Yang, W., Li, A., Mukherjee, A., Basan, M. et al. Protein and lipid mass concentration measurement in tissues by stimulated Raman scattering microscopy. bioRxiv 10.1101/629543, 629543, (2021).

13 Zangle, T. A. & Teitell, M. A. Live-cell mass profiling: an emerging approach in quantitative biophysics. Nat Methods 11, 1221–1228, (2014).

14 Burg, T. P., Godin, M., Knudsen, S. M., Shen, W., Carlson, G., Foster, J. S., Babcock, K. & Manalis, S. R. Weighing of biomolecules, single cells and single nanoparticles in fluid. Nature 446, 1066–1069, (2007).

15 Heimann, P. A. & Urstadt, R. Deep ultraviolet microscope. Applied Optics 29, 495–501, (1990).

16 Barer, R. Determination of dry mass, thickness, solid and water concentration in living cells. Nature 172, 1097–1098, (1953).

17 Zangle, T. A., Chun, J., Zhang, J., Reed, J. & Teitell, M. A. Quantification of biomass and cell motion in human pluripotent stem cell colonies. Biophys J 105, 593–601, (2013).

18 Chun, J., Zangle, T. A., Kolarova, T., Finn, R. S., Teitell, M. A. & Reed, J. Rapidly quantifying drug sensitivity of dispersed and clumped breast cancer cells by mass profiling. Analyst 137, 5495–5498, (2012).

19 Liu, X., Oh, S., Peshkin, L. & Kirschner, M. W. 10.1101/631119, (2019).

20 Reed, J., Chun, J., Zangle, T. A., Kalim, S., Hong, J. S., Pefley, S. E., Zheng, X., Gimzewski, J. K. & Teitell, M. A. Rapid, massively parallel single-cell drug response measurements via live cell interferometry. Biophys J 101, 1025–1031, (2011).

21 <PICS for measuring dry mass changes in sub-cellular compartments.pdf>.

22 Pradeep, S. & Zangle, T. A. Quantitative phase velocimetry measures bulk intracellular transport of cell mass during the cell cycle. Sci Rep 12, 6074, (2022).

23 Bon, P., Maucort, G., Wattellier, B. & Monneret, S. Quadriwave lateral shearing interferometry for quantitative phase microscopy of living cells. Optics Express 17, 13080–13094, p(2009).

24 Barer, R. Interference microscopy and mass determination. Nature 169, 366–367, (1952).

25 Halim, V. A., Garcia-Santisteban, I., Warmerdam, D. O., van den Broek, B., Heck, A. J. R., Mohammed, S. & Medema, R. H. Doxorubicin-induced DNA damage causes extensive ubiquitination of ribosomal proteins associated with a decrease in protein translation. Molecular and Cellular Proteomics 17, 2297–2308, (2018).

26 Kuo, H. K., Chen, Y. H., Wu, P. C., Wu, Y. C., Huang, F., Kuo, C. W., Lo, L. H. & Shiea, J. Attenuated glial reaction in experimental proliferative vitreoretinopathy treated with liposomal doxorubicin. Invest Ophthalmol Vis Sci 53, 3167–3174, (2012).

27 Liaud, N., Horlbeck, M. A., Gilbert, L. A., Gjoni, K., Weissman, J. S. & Cate, J. H. D. Cellular response to small molecules that selectively stall protein synthesis by the ribosome. PLoS Genetics 15, e1008057, (2019).

28 Chen, Y., Hua, Y., Li, X., Arslan, I. M., Zhang, W. & Meng, G. Distinct types of cell death and the implication in diabetic cardiomyopathy. Front Pharmacol 11, 42, (2020).

29 Jung, S., Jeong, H. & Yu, S. W. Autophagy as a decisive process for cell death. Exp Mol Med 52, 921–930, (2020).

30 He, L., Zhang, J., Zhao, J., Ma, N., Kim, S. W., Qiao, S. & Ma, X. Autophagy: the last defense against cellular nutritional stress. Adv Nutr 9, 493–504, (2018).

31 Karim, M. R., Fisher, C. R., Kapphahn, R. J., Polanco, J. R. & Ferrington, D. A. Investigating AKT activation and autophagy in immunoproteasome-deficient retinal cells. PLoS One 15, e0231212, (2020).

32 Edelstein, A., Amodaj, N., Hoover, K., Vale, R. & Stuurman, N. Computer Control of Microscopes Using μManager. 10.1002/0471142727.mb1420s92, (2010).

