## Supplemental Material for "LVING Reveals the Intracellular Structure of Cell Growth"

### Intracellular biomass fixation estimation using LIVING

**The PDF file includes:**

Figs. S1 to S17
Legends for movie M1 to M4
Supplementary derivation of LVING algorithm

**Other Supplementary Material for this manuscript includes the following:**

Movie M1 to M4

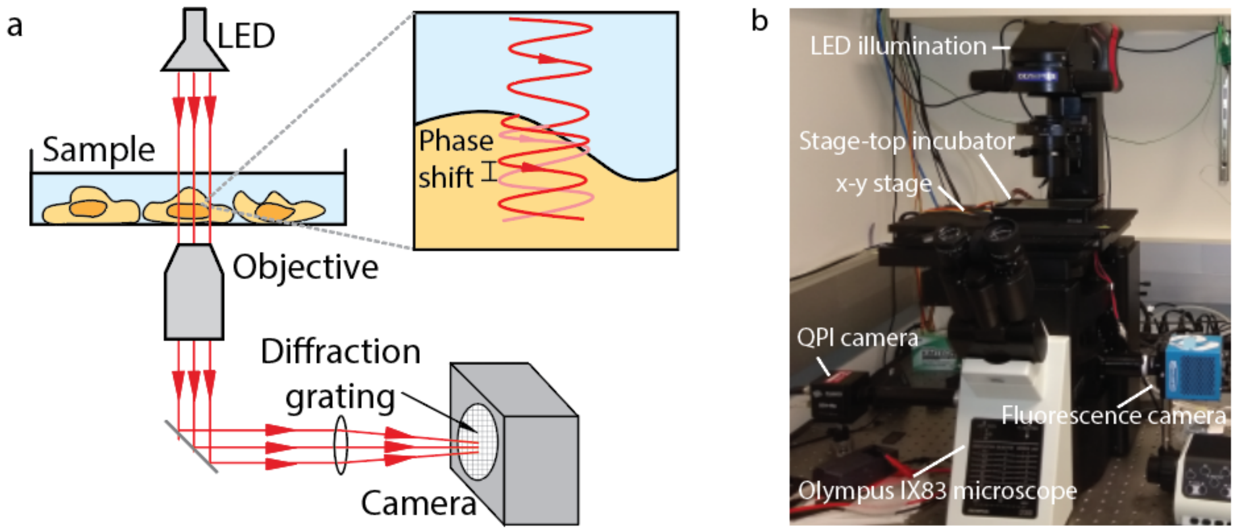


**Figure S1. System used for development of LVING** (a) system diagram, (b) system picture with labels.

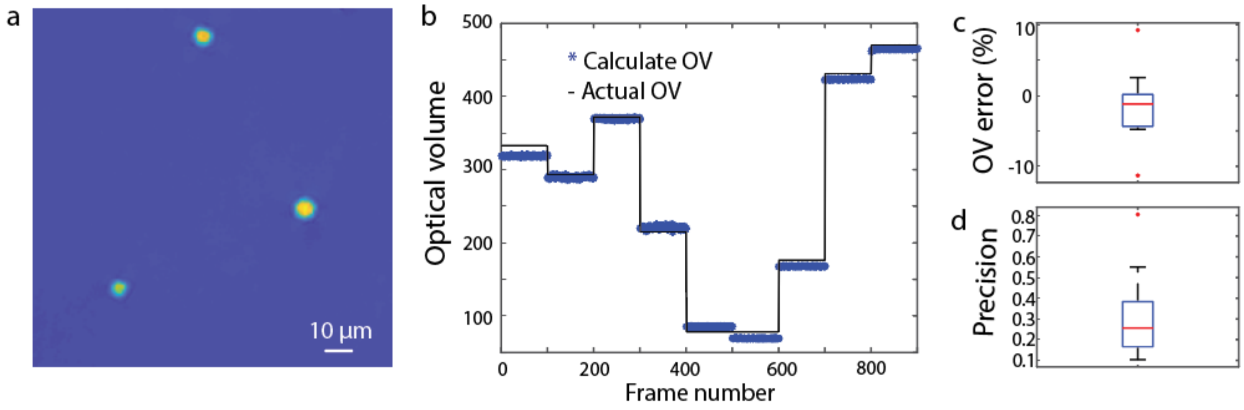


**Figure S2. Verification of QPI system baseline performance** (a) QPI images of polystyrene beads. (b) polystyrene bead optical volume (OV) vs time for 9 imaging locations imaged in sequence. 'Actual' optical volume is based on the known refractive index and measured size of each bead. (c) characterization of error (accuracy) in OV, (d) characterization of precision, measured as the standard error of the mean (SEM) of OV measurements over time.

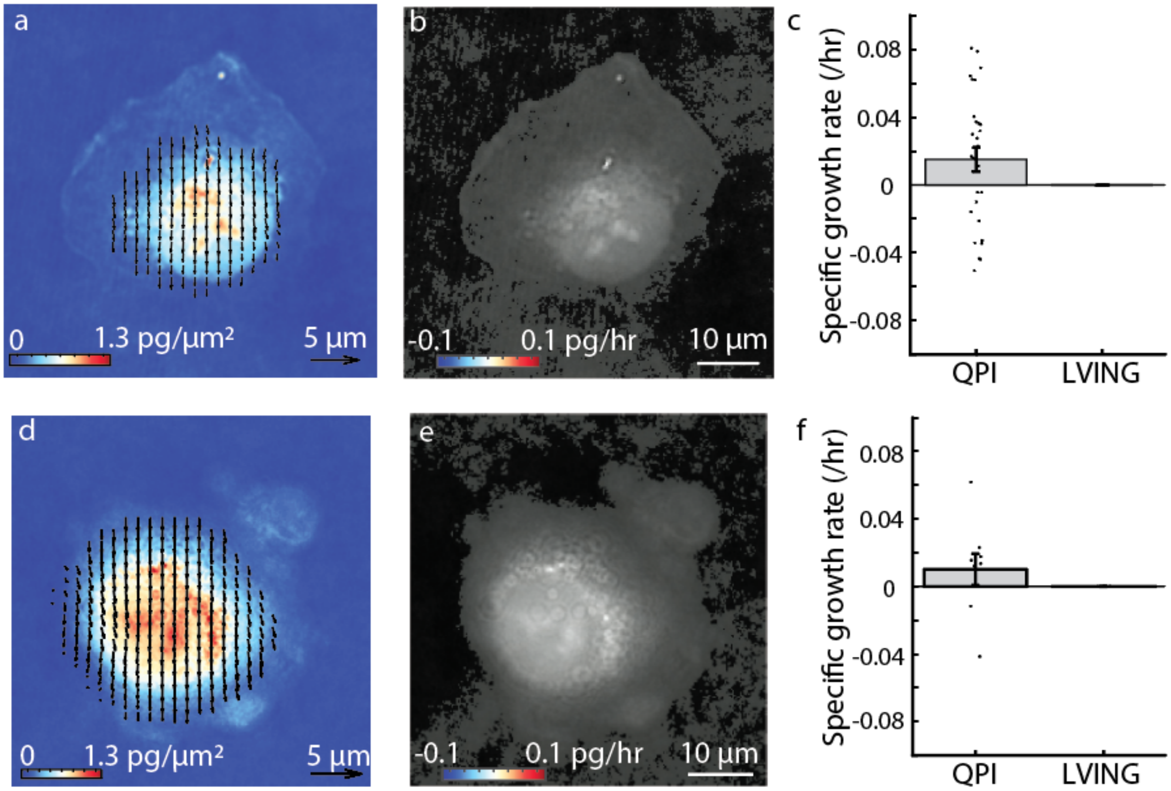


**Figure S3. LVING validation using RPE fixed cells.** (a) Quantitative phase velocimetry (QPV) on representative, fixed RPE cell computes 0.05 μm displacements. (b) Intracellular growth map of the fixed RPE cell shows no growth in any region of the cell, as expected. (c) Specific growth rate computed for $n = 29$ RPE cells from QPI and LVING. **LVING validation using MCF7 fixed cells.** (a) Intracellular transport measurement from QPV shows a representative, fixed MCF7 cell moving at 0.05 μm vertical displacement. (b) Intracellular growth map of the fixed MCF7 shows zero growth, as expected. (c) Specific growth rate computed for $n = 9$ MCF7 cells from QPI and LVING.

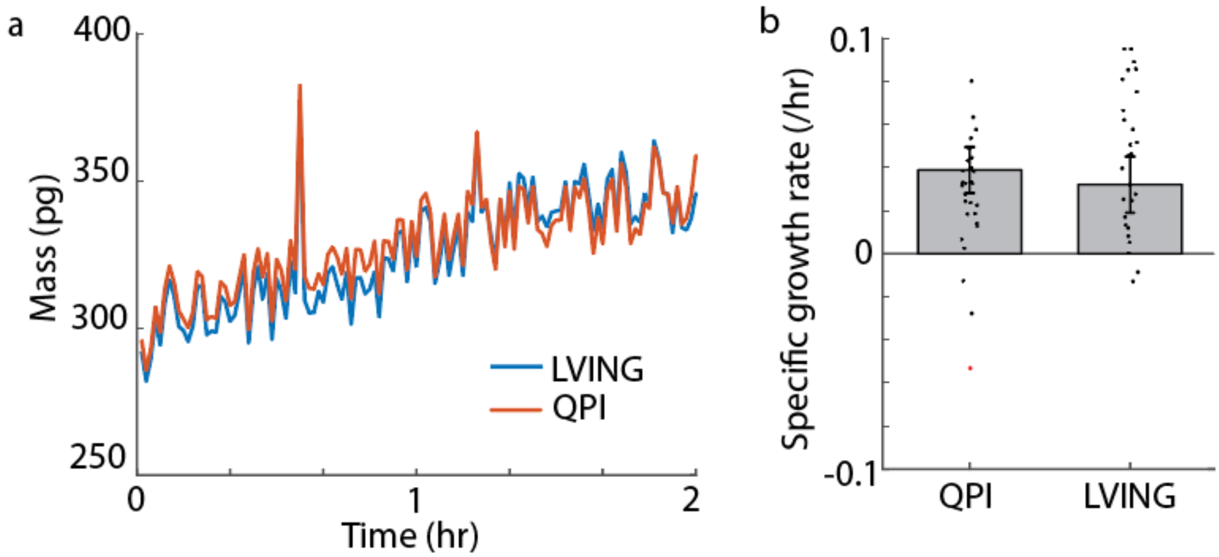


**Figure S4. Comparison of growth measurement using QPI and LVING.** (a) Dry mass vs time for a live cell from QPI and LVING. Blue plot shows the dry mass of the live cell over 2 hours computed from QPI images captured every minute. Red plot shows the sum of dry mass tracks of all 4 by 4 pixel control volumes inside the same cell each Lagrangian tracked over 2 hours. (b) Validation of LVING relative to whole-cell QPI

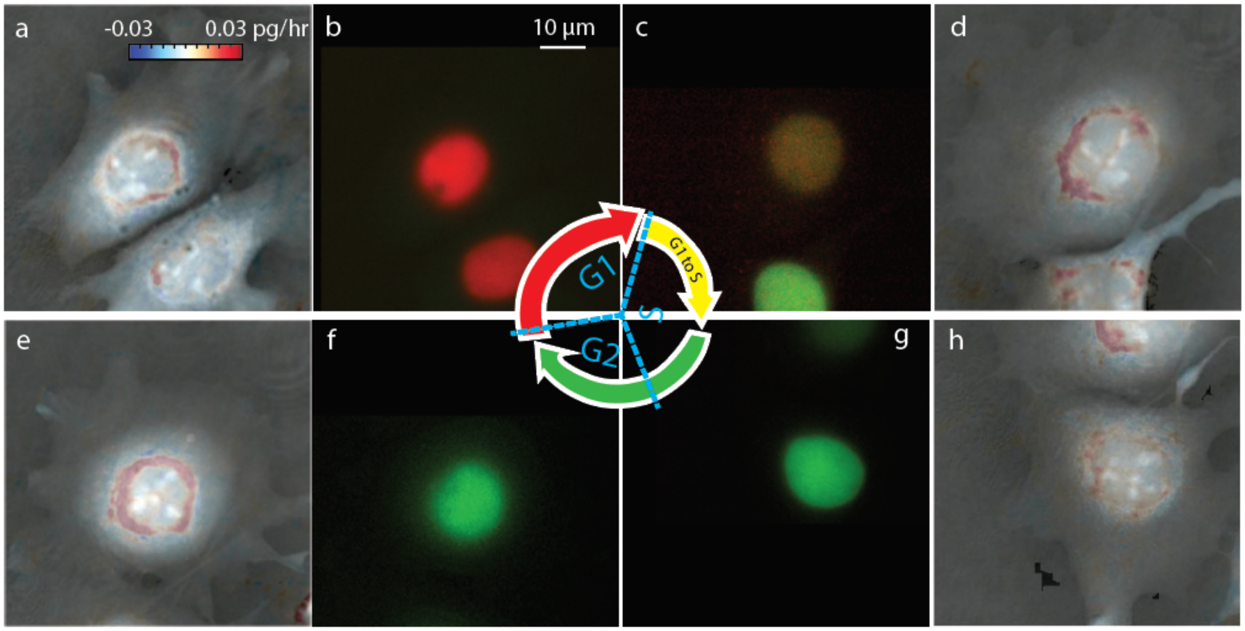


**Figure S5. LVING shows the nuclear and cytoplasm growth in G1, S and G2 phases of the cell cycle in MCF7 cells.** (a) LVING intracellular growth map of MCF7 cell in G1 phase, as indicated by the FUCCI marker in (b) shows a ring of perinuclear growth. (c) The intracellular growth map of the same cells as it progresses to the S phase of cell cycle as indicated by (d) the corresponding FUCCI marker. (d) Growth in perinuclear region of the cell increases as the cells progresses to G2 phase as indicated by (f) the corresponding FUCCI image.

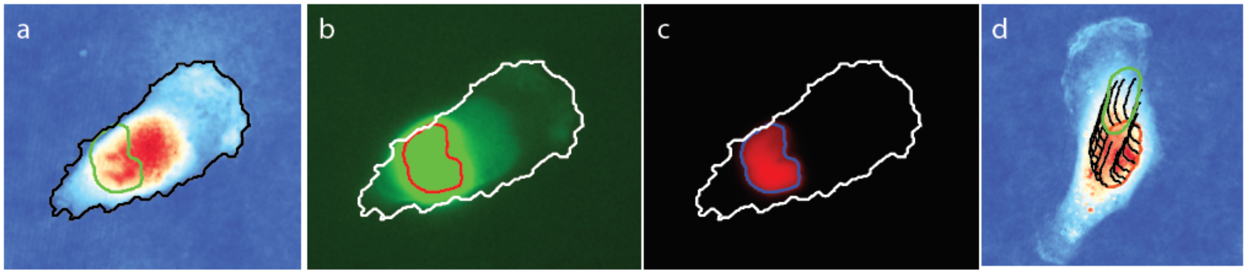


**Figure S6. Subcellular structure tracking using QPV.** (a) nuclear segmentation relative to fluorescence (similar to QPV) (b) Cell nucleus tracked using LVING shows nuclear trajectory without nuclear labelling. The initial position of nuclear boundary shown by purple plot from segmentation of FUCCI label image at the initial time. Dark green plots show the position of handful points selected on the nuclear boundary. Each point is tracked for 30 minutes. Bright green outline shows the position of the nuclear boundary after 30 minutes from FUCCI label image after 30 minutes.

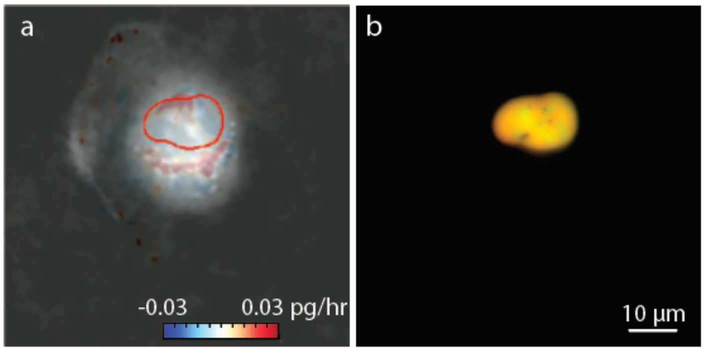


**Figure S7. Growth inhibition by Doxorubicin, a specific inhibitor of DNA synthesis, in RPE cells analyzed by LVING.** (a) production inside the nucleus and cytoplasm at 0.01 µM doxorubicin. Colorbar is growth rate in control volumes in pg/hr. The FUCCI marker in the cells shown in (b) shows that this cell is in the S/G2 phase of cell cycle.

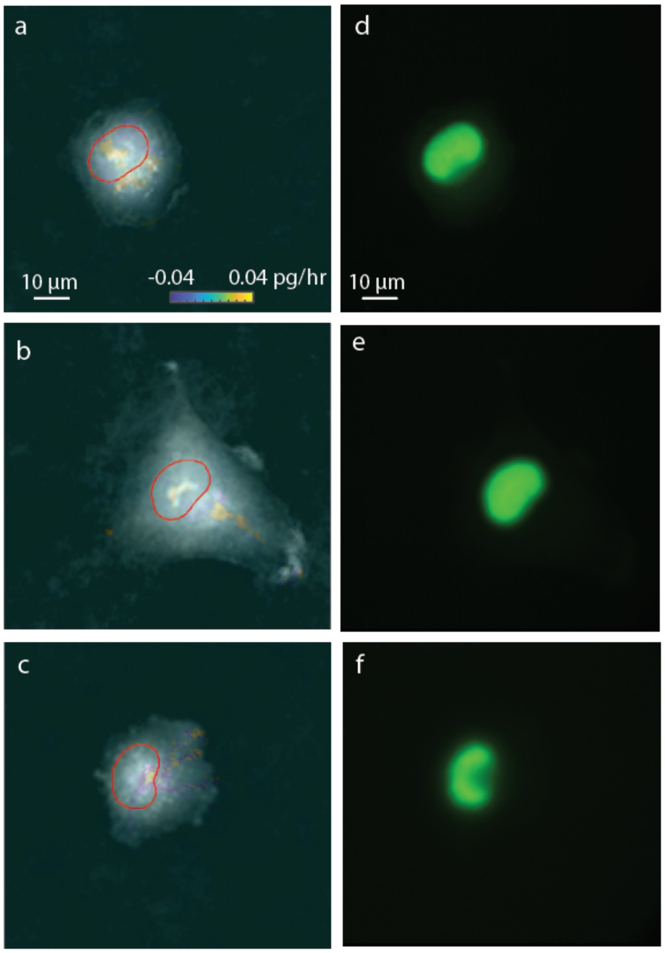


**Figure S8. Growth inhibition by homoharringtonine, a specific inhibitor of protein production, in RPE cells analyzed by LVING.** (a) to (c) Intracellular growth map by LVING indicates biomass production inside the nucleus and cytoplasm at 0.036, 0.36 and 3.6 µM homoharringtonine. Scalebar is 10 µm and colorbar is growth rate in control volumes in pg/hr. The FUCCI marker in the cells shown in (d) to (f) shows that all three cells shown here are in G2 phase of cell cycle.

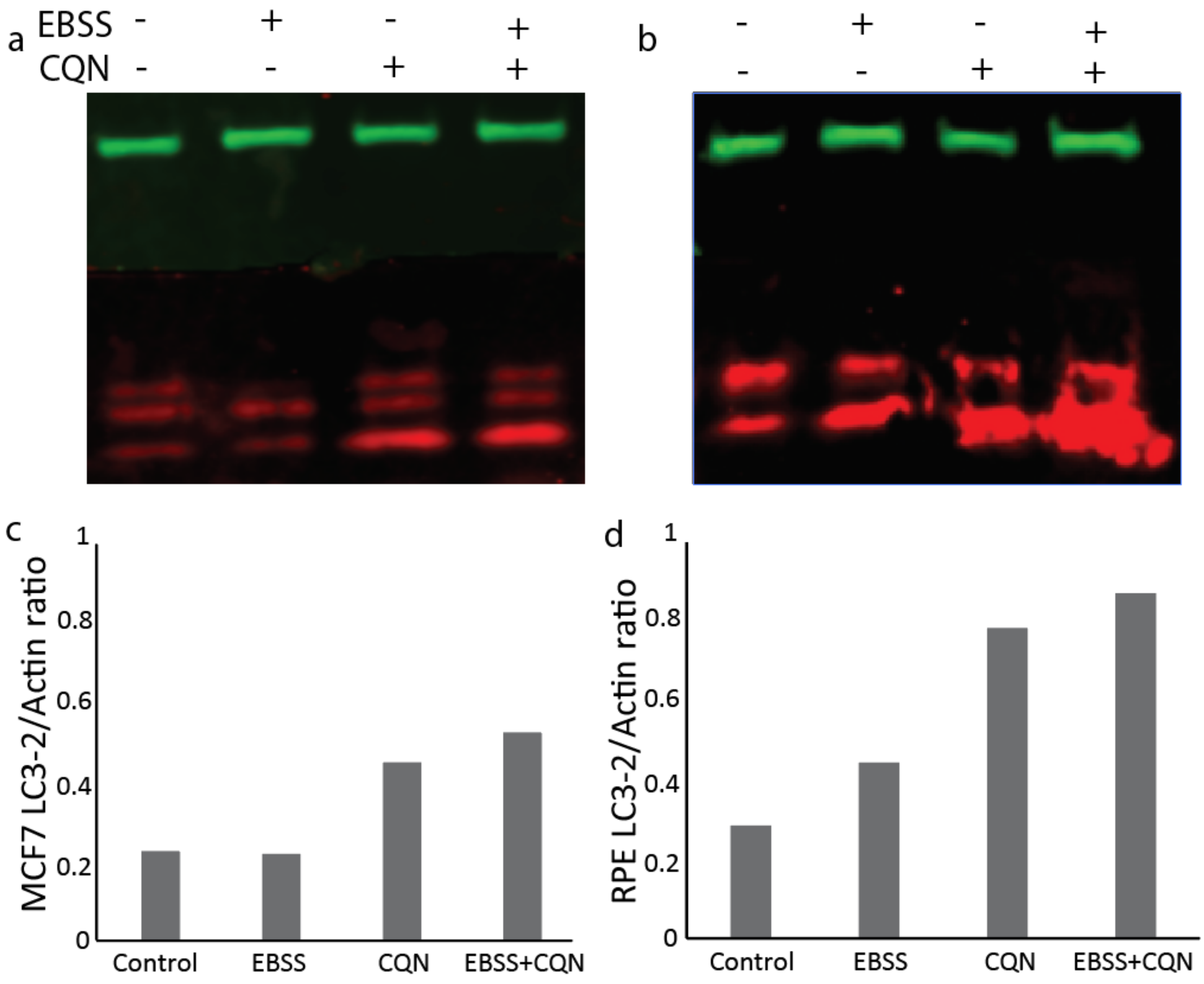


**Figure S9. Western blots and densitometry show MCF7 and RPE cells undergo autophagy in EBBS media.** (a) Western blot shows actin (green), LC3-1 (first red strip) and LC3-2 (bottom-most red strip) for control MCF7 cells (panel 1 vertically), MCF7 cells in EBSS (panel 2), cells with chloroquine (panel 3) and cells in EBSS with chloroquine (panel 4). (b) The densitometry results from MCF7 cells indicates highest LC3-2/Actin ratio in EBSS and chloroquine treated cells as the LC3-2 produced during autophagy is accumulated and degradation is prevented by chloroquine (negative control). (c) Western blot of RPE cells shows actin (green), LC3-1 (top row of red strips) and LC3-2 (bottom thicker row of red strips) for the same conditions as MCF7 cells. (d) Densitometry on the blot indicates increased LC3-2 /Actin ratio in EBSS and chloroquine treated cells similar to MCF7 cells, indicating cells undergo autophagy in EBSS.

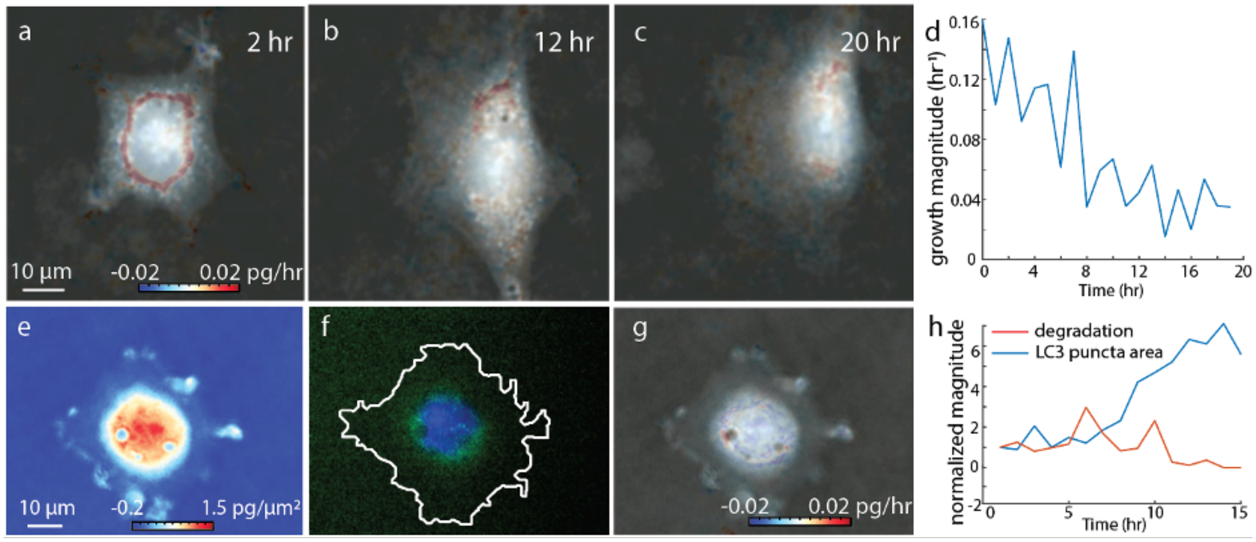


**Figure S10. EBSS induced autophagy in RPE cells shuts down growth observed in cytoplasm.** (a) The intracellular growth map of an RPE cell undergoing autophagy with EBSS treatment at 2, 6, 10, 15 and 20 h shows reducing magnitude of cytoplasmic biomass fixation over time. The nuclear region is marked corresponding to the FUCCI marker using red marker. The scalebar indicates 10μm and colorbar indicates the specific growth rate in control volumes in pg/hr. (b) FUCCI marker indicates the cell observed in (a) is in the G1 phase of cell cycle. (c) The specific growth rate of RPE cells undergoing autophagy over period of 24 h indicates no growth in the cell. The cytoplasmic growth reduces in magnitude over time.

**Movie M1. LVING data from RPE cell through the cell cycle.** Left: LVING growth rate map superimposed on gray scale QPI mass distribution. Right: FUCCI cell cycle marker indicating cell cycle phase.

**Movie M2. LVING data from MCF7 cell through the cell cycle.** Left: LVING growth rate map superimposed on gray scale QPI mass distribution. Right: FUCCI cell cycle marker indicating cell cycle phase.

**Movie M3. LVING data from RPE cell undergoing autophagy.** Left: LVING growth rate map superimposed on gray scale QPI mass distribution. Right: FUCCI cell cycle marker indicating cell cycle phase.

**Movie M4. LVING data from MCF7 cell undergoing autophagy.** Left: LVING growth rate map superimposed on gray scale QPI mass distribution. Right: FUCCI cell cycle marker indicating cell cycle phase.

**Supplementary Derivation: Eulerian to Lagrangian transformation for LVING**

***Statement of mass conservation***

The basic Eulerian form of mass conservation, allowing for the generation of mass as biosynthesis, states that:

$$\begin{Bmatrix} rate\ of \\ increase \\ of\ mass \end{Bmatrix} = \begin{Bmatrix} rate\ of \\ mass\ in \end{Bmatrix} - \begin{Bmatrix} rate\ of \\ mass\ out \end{Bmatrix} + \begin{Bmatrix} rate\ of \\ mass \\ generation \end{Bmatrix} \tag{1}$$

For a cuboid control volume of sides Δx, Δy and Δz:

$$\Delta x \Delta y \Delta z \frac{\partial \rho}{\partial t} = \Delta y \Delta z[(\rho v_x)|_x - (\rho v_x)|_{x+\Delta x}] + \Delta \mathrm{x} \Delta \mathrm{z}\left[(\rho v_y)|_y - (\rho v_y)|_{y+\Delta \mathrm{y}}\right] + \Delta \mathrm{x} \Delta \mathrm{y}[(\rho v_z)|_z - (\rho v_z)|_{z+\Delta \mathrm{z}}] + r \tag{2}$$

where ρ is the mass density, and *r* is the rate of biomass production. This gives:

$$\frac{\partial \rho}{\partial t} = \left[\frac{1}{\Delta x}(\rho v_x)|_x - (\rho v_x)|_{x+\Delta x}\right] + \frac{1}{\Delta \mathrm{y}}\left[(\rho v_y)|_y - (\rho v_y)|_{y+\Delta \mathrm{y}}\right] + \frac{1}{\Delta \mathrm{z}}[(\rho v_z)|_z - (\rho v_z)|_{z+\Delta \mathrm{z}}] + \frac{r}{\Delta \mathrm{x} \Delta \mathrm{y} \Delta \mathrm{z}} \tag{3}$$

Simplifying and applying the del operator (∇) yields:

$$\frac{\partial \rho}{\partial t} + \nabla \cdot \rho v = \frac{r}{\Delta V} \tag{4}$$

Multiplying through by the volume, $\Delta V$:

$$\frac{\partial m}{\partial t} + \nabla \cdot m v = r \tag{5}$$

In Lagrangian framework eq (5) can be written as,

$$\frac{Dm}{Dt} + v \cdot \nabla m = r \tag{6}$$

The total derivative is calculated since the tracking is performed in Eulerian framework accounting for the change in mass while the CV is in motion. Partial derivative is used when tracking performed in Eulerian framework. The total derivative $\frac{Dm}{Dt}$ $is\ defined\ as$:

$$\frac{Dm}{Dt} = \frac{\partial m}{\partial t} + m\nabla \cdot v \tag{7}$$

Expanding,

$$\frac{Dm}{Dt} + v_x \frac{\partial m}{\partial x} + v_y \frac{\partial m}{\partial y} + v_z \frac{\partial m}{\partial z} = r \tag{8}$$

Assuming minimal change in height we assume $v_z << v_x$ and $v_y$:

$$\frac{Dm}{Dt} + v_x \frac{\partial m}{\partial x} + v_y \frac{\partial m}{\partial y} = r \tag{9}$$

Here, the gradient of mass across x and y direction in individual CVs are negligible as we track the smallest CV at the highest time resolution (**figure S18**).

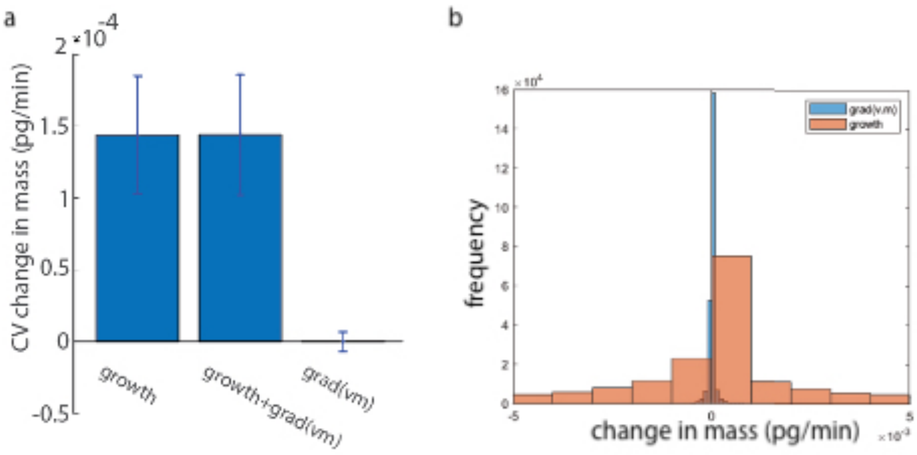


**Figure S18. Comparison of terms from LVING derivation indicates that grad(*vm*) term is negligible.** (a) comparison of average growth ($r$), growth + grad($vm$), and grad($vm$). (b) distribution of control volume growth ($r$) and computed grad($vm$).

***Lagrangian to Eulerian transformation for biomass production measurement***

The first term in eq 9 indicates the change in mass during translation of a fixed control volume over time. Overall, eq. 9 states that mass inside a fixed control volume can change with processes like protein production (*r*) as well as deformation of the CV ($v_x \frac{\partial m}{\partial x} + v_y \frac{\partial m}{\partial y}$), with other terms negligible. Therefore, to account for this, and recover the biomass production rate, *r*, we must translate from the Eulerian statement of mass conservation (eq. 9) to one that accounts for deformation of the control volumes within the cell over time.

We start by considering the net mass generation, $m_{gen}$ within a Lagrangian CV that deforms with overall cellular motion:

$$m_{gen} = m_t - m_0 \tag{10}$$

Where $m_t$ is the mass inside the CV after the translation and deformation and $m_0$ is the mass in the CV at initial time point. We can obtain the rate of change of mass with time due to intracellular activity as:

$$\frac{m_{gen}}{\Delta t} = r \tag{11}$$

When extended to multiple measurements of a deforming CV over time, the rate of generation term, *r*, is, therefore, approximately the linear slope of mass of the deforming CV as a function of time.

To obtain mass as a function of time for each deforming CV, we use computed velocity vectors for intracellular mass transport[1] to track the corners of quadrilateral CVs (**figure S19**). We, therefore, assume that the sides of the control volume do not bend with intracellular deformations. This assumption is valid in the limit of small CVs. We then update the position of the coordinates at the corners of the CV at each time step as PIV progresses.

Phase shift is measured at each pixel in Eulerian coordinates on a fixed grid. In order to calculate the mass inside the CV at each time step we use the shoelace algorithm to measure the area of CV, which is area inside a quadrilateral overlayed on a grid, then multiply by the average phase shift at that location. As an example of implementation of the shoelace algorithm, we show the area inside a square of side xcg and further a non-uniform side quadrilateral as in **figure S19**.

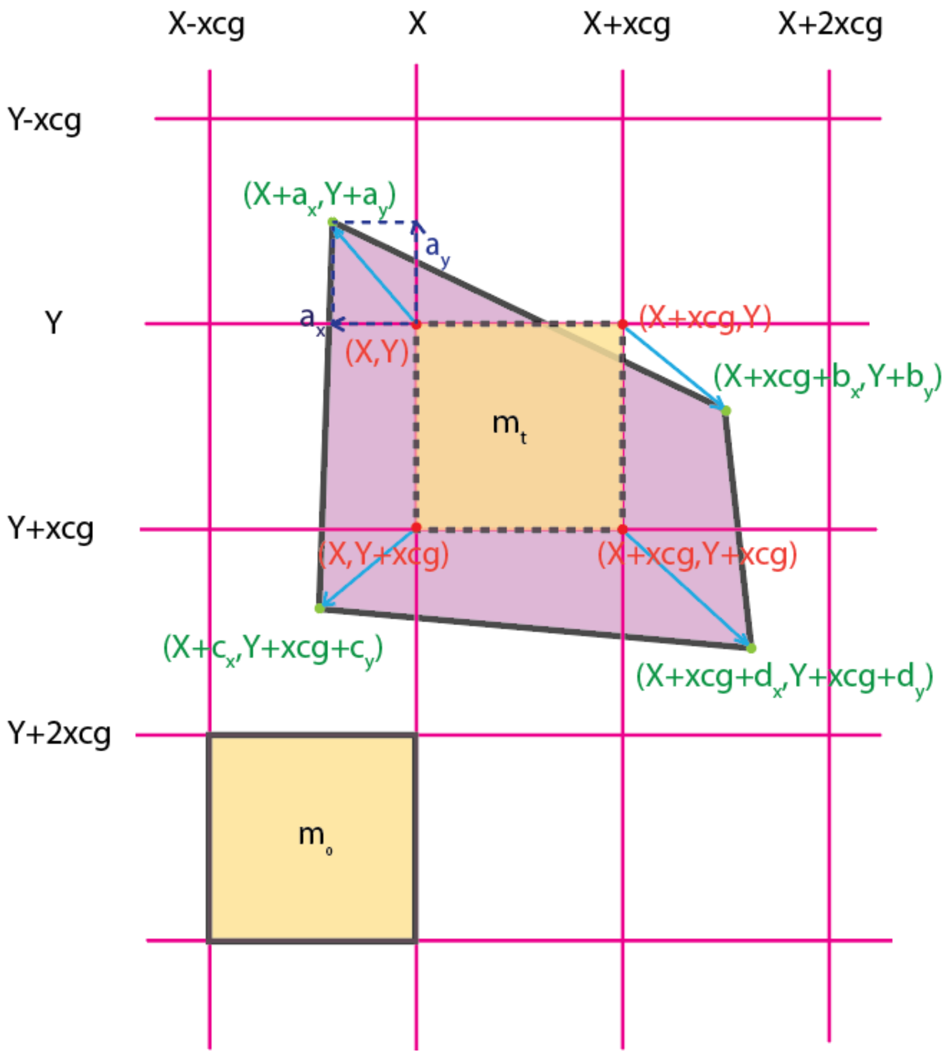


**Figure S19. Deformation of quadrilateral control volume tracked by LVING.** Square control volume of mass $m_0$ deform over time $\Delta t$ to a quadrilateral control volume of mass $m_t$. The corners of the control volume are displaced due to overall translation as well as control volume deformation.

For a square of sides xcg, overlayed on a grid of origin (X,Y), using the shoelace algorithm we obtain,

$$Area1 = \left|\frac{1}{2}[X(Y + xcg) + X(Y + xcg) + Y(X + xcg) + Y(X + xcg) - 2XY - 2(X + xcg)(Y + xcg)]\right| = xcg^2 \quad (12)$$

If phase shift inside the square CV is $\phi_l$, dry mass ($m_O$) inside the CV is,

$$m_t = \phi_t \times xcg^2 \quad (13)$$

At the next time point the CV volume can be a non-uniform side quadrilateral due to deformation and translation of the cell. For a non-uniform side quadrilateral overlayed on the same grid as above as shown in **figure S19**, we can perform the area calculation using the shoelace algorithm.

$$\begin{aligned} Area2 = \Big|\frac{1}{2}&[(X + a_x)(Y + xcg + C_y) + (X + c_x)(Y + xcg + d_y) \\ &+ (X + xcg + d_x)(Y + b_y) + (X + xcg + b_x)(Y + a_y)] \\ &- \frac{1}{2}[(Y + a_y)(X + c_x) + (Y + xcg + c_y)(X + xcg + d_x) \\ &+ (Y + xcg + d_y)(X + xcg + b_x) + (Y + b_y)(X + a_x)]\Big| \\ &= \Big|\frac{1}{2}[(a_x - xcg)(c_y - b_y) + (c_x - xcg)(d_y - a_y) + d_x(b_y - c_y) \\ &+ b_x(a_y - d_y) + xcg(a_x + c_x - d_x - b_x)] - xcg^2\Big| \end{aligned} \quad (14)$$

Dry mass in the CV at time $t+\Delta t$, $m_{t+\Delta t}$, is product of phase shift ($\phi_{t+\Delta t}$) and area (*Area2*) at each pixel. In order to account for this, we use the shoelace algorithm to compute a mask with pixel values representing the total area contribution of each pixel to the deformed control volume. This is multiplied by the phase shift to obtain $m_{t+\Delta t}$.

Substituting $m_t$ and $m_{t+\Delta t}$ into equation 11, we can calculate the instantaneous growth rate between the two time points. However, this approach is often noisy. Therefore, LVING calculates the slope of the curve of mass in CV vs time to estimate *r*, the rate of growth at that control volume.

1 Pradeep, S. & Zangle, T. A. Quantitative phase velocimetry measures bulk intracellular transport of cell mass during the cell cycle. *Sci Rep* **12**, 6074, (2022).
